# A VHL-1;HIF-1/SQRD1/COL-88 axis links extracellular matrix formation with longevity in *Caenorhabditis elegans*

**DOI:** 10.1101/2024.02.22.581513

**Authors:** Willian Salgueiro, Reza Esmaillie, Katrin Bohl, Cyril Statzer, Puneet Bharill, Sebastian Bargfrede, Manopriya Chokkalingam, Maike Neutzer, Michael Ignarski, Thomas Benzing, Andreas Beyer, Bernhard Schermer, Collin Y. Ewald, Francesca Fabretti, Roman-Ulrich Müller

## Abstract

The extracellular matrix (ECM) is a pivotal three-dimensional network crucial for tissue organization, cellular communication, and fundamental cellular processes, where collagens are the major chemical entity in amount. ECM deregulation is directly involved with several pathologies, such as tumour growth and invasiveness, atherosclerosis, and diabetic nephropathy. Mutations in the von Hippel-Lindau tumour suppressor (pVHL) cause VHL syndrome, a multi-tumour syndrome commonly associated with clear cell renal carcinoma (ccRCC). Loss of pVHL is associated with the activation of hypoxia-inducible factor (HIF) signaling. Mutation of VHL-1 in the nematode *Canorhabditis elegans* has been shown to increase lifespan and stress resistance. Interestingly, considering recent findings on the involvement of collagens in the regulation of lifespan, we also observed these animals to show defects in body morphology in a HIF-1 dependent manner. Based on this finding, we established a link between HIF-1 activation upon loss of VHL-1 and ECM defects associated with alterations in collagen expression. An RNAi screen examining genes upregulated in *vhl-1* mutant worms revealed the sulfide quinone oxidoreductase *sqrd-1* to mediate the change in body morphology. SQRD-1 is essential to the HIF-1 dependent increase in several collagen genes. One of these genes, *col-88*, partly mediates both the impact of loss of VHL-1 on lifespan extension and body length. The downregulation of the uncharacterised *col-88* partially restores lifespan extension and reduces body size of *vhl-1/sqrd-1* to *vhl-1(ok161)* single mutant. This study contributes to the increasing body of evidence linking lifespan extension and the ECM and now implicates this axis in hypoxia-signaling. These findings are of special interest considering the role of ECM integrity in tumour growth and metastasis.

**Author Summary:** The extracellular matrix and its composing collagens are associated with a wide number of diseases, including cancer. The von Hippel-Lindau tumour suppressor (pVHL) is known to work by regulating the Hypoxia Inducible Factor (HIF) to help the organism to adapt to lack of oxygen. Mutations in pVHL are associated with clear cell renal carcinoma (ccRCC). Interestingly, a small number of studies have shown that pVHL can be directly associated with collagens, a function that is independent of its classical role regulating HIF. However, there is no further knowledge about which role the hypoxia pathway has when it comes to extracellular matrix formation and function, what would be useful since the invasiveness of cancers, such as ccRCC, are directly connected to their matrix/collagen composition. Here we observed that the model organism *C. elegans* has drastically different collagen composition and body size upon a mutation on its *vhl-1* gene. Furthermore, a protein previously only known to be involved in sulfide metabolism, SQRD-1, connects body size and lifespan in this animal model, revealing a surprising link between the hypoxia pathway and sulfur metabolism to control lifespan. Further studies could target sulfur metabolism in ccRCC to modulate collagen production and tumour invasiveness.

## Introduction

The von Hippel-Lindau tumour suppressor (pVHL) tightly regulates the Hypoxia Inducible Factor (HIF) across phylae (1,2), and HIF regulates an orchestrated expression of a plethora of genes, so that the organism can properly adapt to hypoxic conditions (1). In this regulatory network, pVHL is the substrate recognition subunit of an E3 ubiquitin ligase mediating the proteasomal degradation of the HIF alpha subunit (3,4). Much is known about how pVHL interaction with HIF-1 contributes to tumorigenesis through induction of genes known to be relevant for tumor cell growth, e.g. VEGF, TGFα, and PDGFβ (5–8). However, pVHL has also been shown to have an impact on extracellular matrix (ECM) formation and composition, both in a HIF-1 dependent and -independent manner (9–12) (13,14). This is of interest, since ECM modulation has important implications in tumor invasion and metastasis. Interestingly, ECM homeostasis seems to be linked to the aging process and anti-aging interventions as well (15). Besides, in the nematode *Caenorhabditis elegans (C. elegans)*, HIF-1 activation through loss of the nematode orthologue of pVHL (VHL-1) has been shown to induce longevity(16,17). ECM related genes have been shown to be regulated not only by pVHL/VHL-1 (18)several other well-established longevity-related transcription factors (*e.g.* FOXO3, Nrf2, TP53) (15). Furthermore, established longevity inducing compounds such Rapamycin, Metformin, and Aspirin do also change ECM composition in a number of model organisms (19) Most importantly, ECM modulation has been linked directly and causally to lifespan-extension in *C. elegans* (20). Interestingly, we had observed a body morphology phenotype upon VHL-1 knockout implicating a role for VHL-1/HIF-1 in *C. elegans* ECM formation. Combining targeted genetics with screening approaches we identified a novel VHL-1/HIF-1 axis which – through induction of the Mitochondrial Sulfide Quinone Oxidoreductase SQRD-1 – unifies ECM and longevity-associated phenotypes in a collagen-dependent manner.

## Results

### *vhl-1(ok161)* shows a *hif-1* dependent phenotype in body length

When working with *vhl-1(ok161)* mutants, we repeatedly observed an abnormal body morphology throughout different larval stages, when compared to wild type N2 (WT). We confirmed this finding by applying the WormSizer plugin (21) in ImageJ, which showed significant differences in body length between *vhl-1(ok161)* mutants and WT at the larval stage 1 (L1) (**Fig 1A/B**). Considering the crucial role of VHL-1 in the degradation of hypoxia-inducible factors, we examined if this phenotype depended *on hif-1*. Regarding morphology, *hif-(ia4)* (251µm) mutants did not show an overt phenotype. Importantly, while *vhl-1(ok161)* L1 larvae (187µm) are 84µm shorter than WT worms (271µm), the double mutant *vhl-1;hif-1* (236µm) is indistinguishable from WT worms (**Fig 1B**). The length phenotype was also easily detectable in L4 larvae and was again rescued by loss of *hif-1* (**Fig 1C**). The phenotype was limited to differences in length; width of both L1 and L4 larvae showed no significant differences (**Suppl. Fig. 1A and 1B**). We went on to test whether the phenotype would have an impact on the formation of dauer larvae, a highly resilient larval stage of the nematode which develops upon stress. *vhl-1(ok161)* dauers indeed were resistant to 1% SDS after starvation (**Suppl. Fig. 1C**), even though a smaller number of *vhl-1(ok161)* animals would go into the dauer stage when compared to WT. However, again in a *hif-1* dependent manner, these animals differed morphologically from a regular dauer animal and were shorter than WT dauers (**Fig. 1D** and **E**). Taken together, loss of *vhl-1* induces a *hif-1* mediated body morphology phenotype across different developmental stages of the nematode.

**Figure 1.**
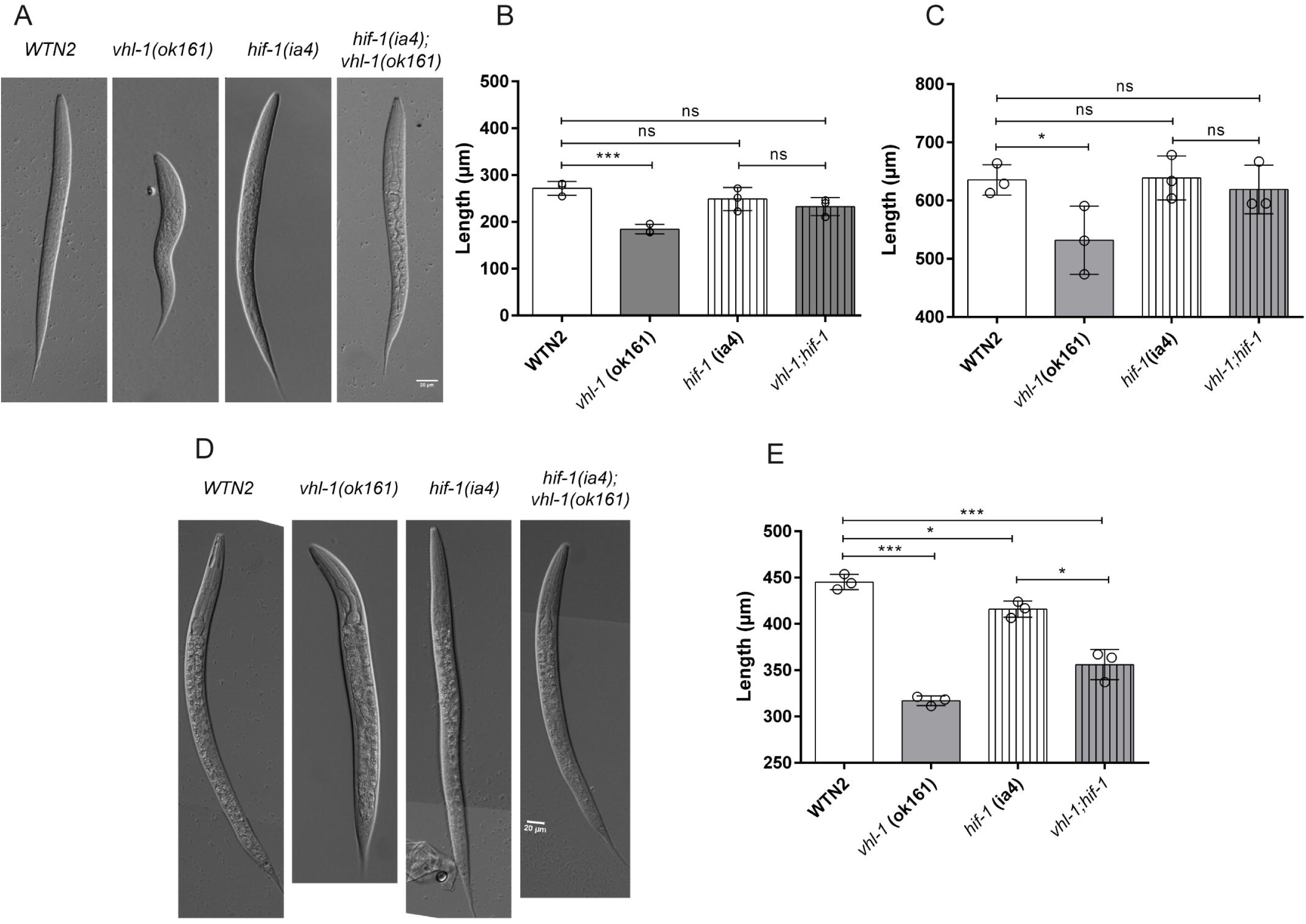
*vhl-1(ok161)* mutation shortens *C. elegans* body in a *hif-1* dependent manner. **A)** Representative images of L1 WT worms, *vhl-(ok161)*, *hif-(ia4)* and *hif-(ia4)*;*vhl-(ok161)*. The scale bar represents 20µm; **B)** Length of L1 worms; **C)** Length of L4 worms; **D)** Representative images of dauer stage WT *vhl-(ok161)*, *hif-1(ia4)* and *hif-1(ia4)*;*vhl-1(ok161)*; **E)** Length of dauer worms; Each point reflects the length of a single worm. The three dots represent the mean of three independent experiments, while the black line represents the mean of these three experiments. Significance was calculated by comparing the mean value of the three independent experiments by a One-Way ANOVA followed by a Tukey post-hoc. ns = not significant, *** = p<0.001, * = p<0.05.

### Transcriptome analyses link HIF-signaling to changes in collagens and other genes involved in the regulation of extracellular matrix formation

To identify potential mediators contributing to the shorter size observed in *vhl-1(ok161)* mutants, we acquired RNAseq data from these mutants compared to N2, as well as, from *hif-1(ia4)* and the *vhl-1;hif-1* double mutants. **Fig 2A** displays the Z score transformed gene expression per sample for all regulated genes (as by ANOVA and a p<0.05). Hierarchical clustering separated *vhl-1* from both WT and *vhl-1;hif-1* worms. The analysis resulted in three clusters of genes, two of which (red and blue) appeared to be clearly *hif-1* dependent.

**Figure 2.**
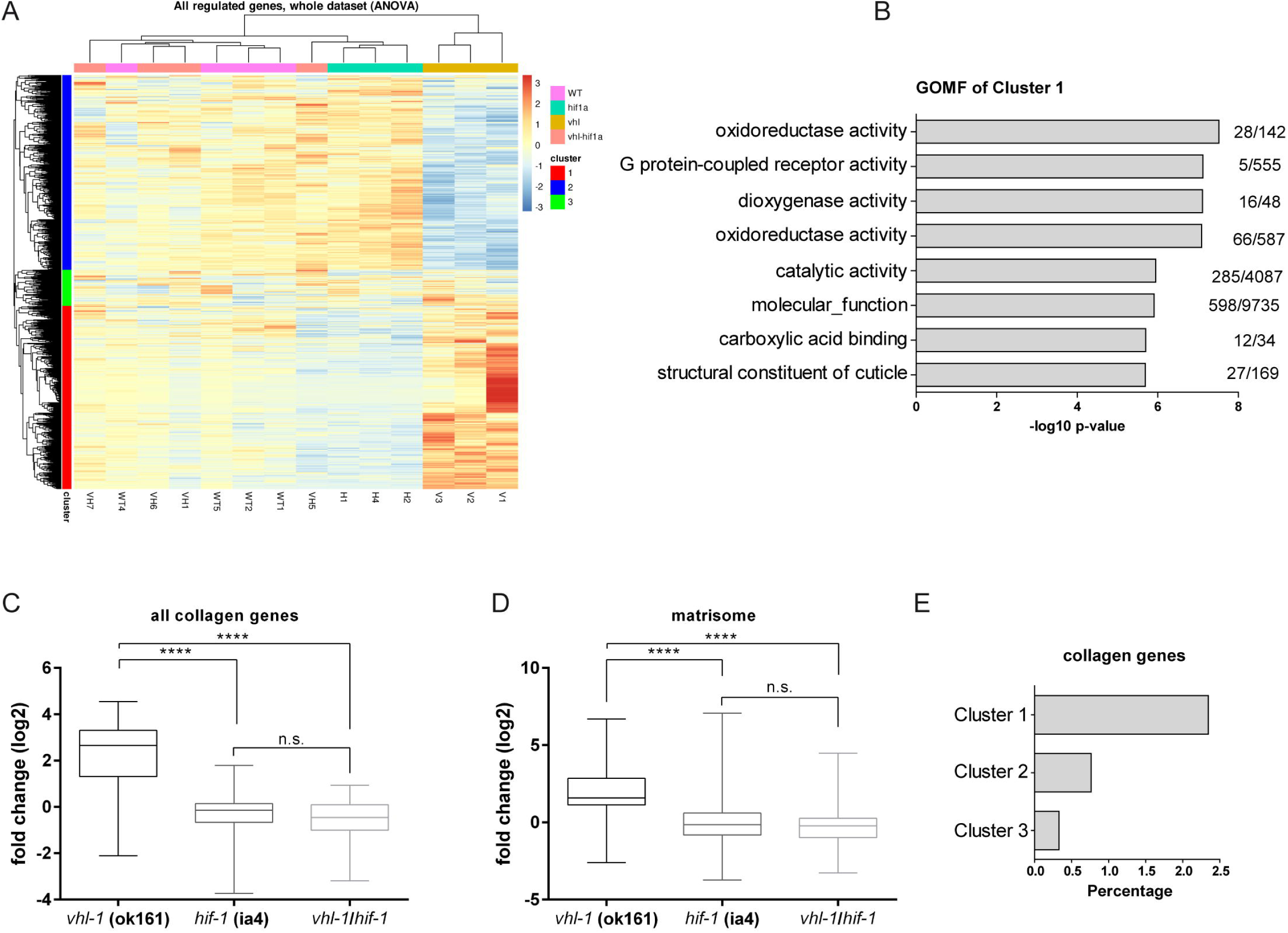
RNAseq data reveals a global change in the expression of collagen genes. **A)** RNAseq heatmap. The Z score transformed gene expression per sample for all differentially regulated genes (ANOVA, adj. p-value < 0.05) is displayed. Columns are samples, rows are genes. The first cluster, depicted in red, depicts genes that are highly expressed in *vhl-1(ok161)* mutant; **B)** Gene Ontology Molecular Function (GOMF) of RNAseq data. Top eight GO-terms with the highest enrichment factor. Numbers indicate proteins in the category followed by category size; **C)** Log2 fold change of all *C. elegans* collagen genes detected in our RNAseq data for each genotype; **D)** Log2 fold change of extracellular matrix components (matrisome (22,23)) genes detected in our RNAseq data for each genotype; **E)** Collagen genes distribution of clusters 1-3. Significance was calculated by comparing the mean value of the three independent experiments by a One-Way ANOVA followed by a Tukey post-hoc. ns = not significant, **** = p<0.0001

Interestingly, in a Gene Ontology analysis for Molecular Function in Cluster 1 containing genes which were upregulated upon loss of *vhl-1* in a *hif-1* dependent manner (**Fig 2B**), “structural constituent of cuticle” was among the top 8 enriched GO-terms. We examined the expression pattern of collagen-encoding genes in *vhl-1(ok161)* mutants. We observed that *vhl-1(ok161)* mutants have an overall upregulation of all collagen genes on average, which is reversed by addition of *hif-1(ia4)* mutation (**Fig. 2C)**. Collagen genes are mostly represented in Cluster 1 (**Fig. 2E**), where they sum over 2/3 of all collagen genes detected when compared to the whole dataset. Also, all genes that fall under the umbrella of extracellular matrix components (including collagens, but also proteoglycans and glycoproteins (22)), the matrisome, were regulated in a similar manner (**Fig. 2D**) and genes associated with a dumpy phenotype (22) showed a similar pattern (**Suppl. Fig. 1A**). By contrast, this was not the case for genes associated with other ECM phenotypes such as ‘’Long (lon)’’ and ‘’Roller (rol)’’, which are defined by an elongated body and a helically twisted body, respectively (**Suppl. Fig. 1B and C**). In conclusion, loss of *vhl-1* induces *hif-1* dependent changes in ECM-associated transcript abundance.

### Extracellular matrix composition and functionality is altered in *vhl-1(ok161)* mutants

To confirm whether the increased collagen gene expression changes in the profile of expression of overall collagen genes would impact total collagen on the protein level in *C. elegans vhl-1(ok161)* mutants, we measured total collagen by colorimetric quantification of hydroxyproline, a relatively specific marker due to the abundant common proline hydroxylation events in collagens (24) Adult *vhl-1(ok161)* mutants showed higher levels of hydroxyproline than WT (**Fig. 3A**) Consistent with that worms with a mutation in the insulin/IGF-1 receptor *daf-2*, for which an increase in collagen levels had previously been shown (20), were used as a positive control (**Fig. 3A**). Furthermore, a highly expressed and well-established collagen reporter, *col-19*p::*col-19*::GFP, is more abundant at protein levels in day 8 adult (D8) *vhl-1(ok161)* mutants (**Fig. 3B** and **3C**), while D1 adults did not show a change in the levels of this fusion protein presumably since the *col-19* expression starts at the transition from the last larval stage L4 to D1 adults (**Suppl. Fig 2A and B**).

**Figure 3:**
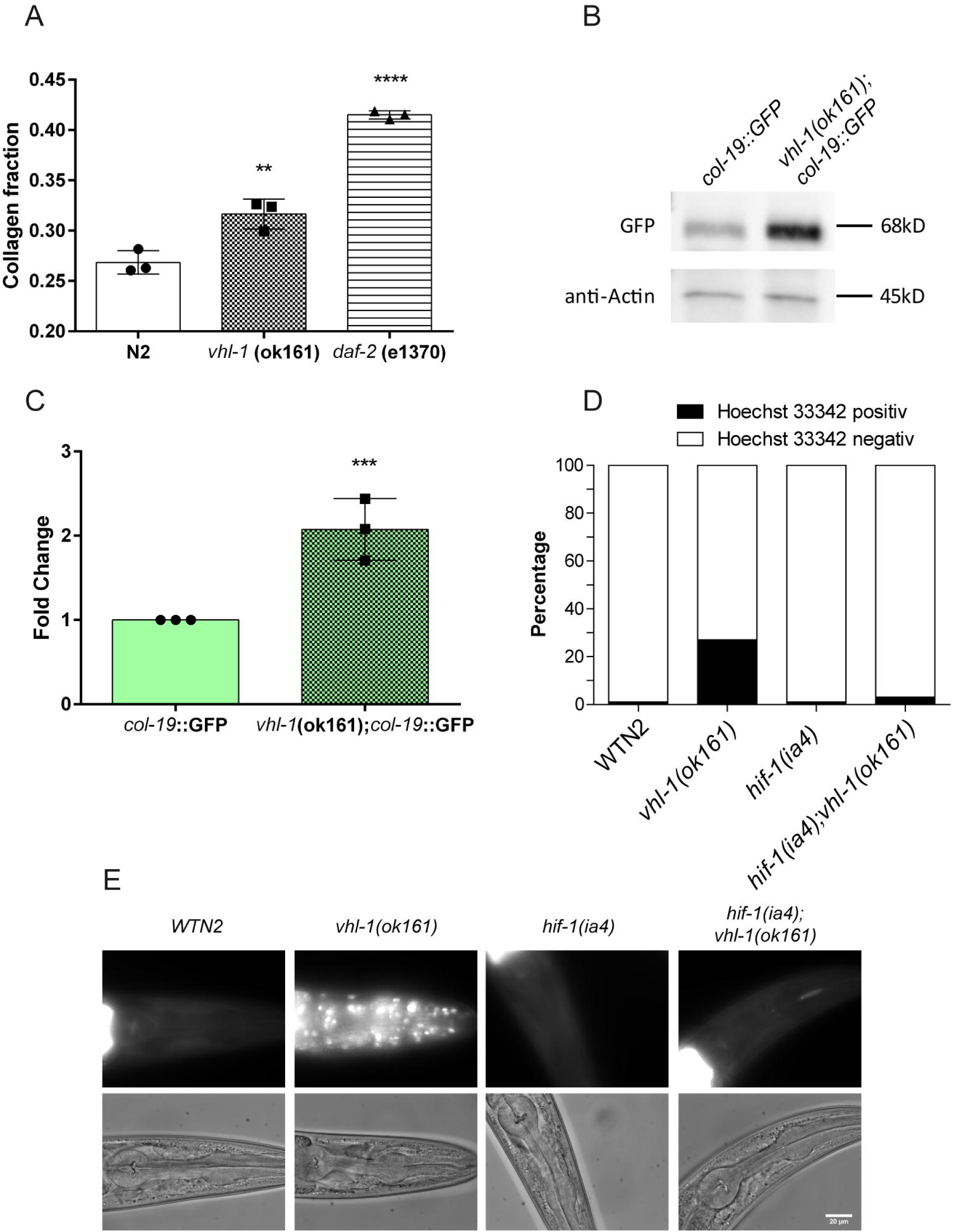
*vhl-1(ok161)* mutants ECM is functionally impaired. **A)** Hydroxyproline colorimetric measurement in day one old WT, *vhl-1(ok161)*, and *daf-2* (e1370); **B** and **C)** Representative Western blot images and densitometry measurements of anti-GFP stainings of *col-19*p*::col-19::*GFP reporter in WT and *vhl-1(ok161)* backgrounds in day 8 old adult worms; **D** and **E)** Fluorescence measurement and representative images of Hoechst diffusion into the cuticle of different genotypes in D1 worms; Significance was calculated by comparing the mean value of the three independent experiments by an One-Way ANOVA followed by a Tukey post-hoc. * = p<0.05, ** = p<0.01, *** = p<0.001.

We continued to assess the impact of these gene expression changes on cuticle functionality, the primary component of nematodes made up by collagen and determining body shape (25). This was examined using Hoechst staining, a dye which is membrane permeable but does usually not permeate the cuticle (26). In contrast to WT, *vhl-1(ok161)* animals displayed abnormal Hoechst positivity in the head (**Fig. 3D** and **E**), suggesting that the cuticle is defective [25,26]. Again, loss of *hif-1* fully rescued the phenotype of *vhl-1* mutant worms and *hif-1(ia4)* single mutants did not show differences to WT. It is interesting to notice that our *vhl-1(ok161)* mutants are long-lived and heat stress resistant (17), but we have recently accessed their stress resistance against paraquat and observed that indeed they are less resistance to that toxicant (**Suppl. Fig. 2C**). To address the question whether loss of *vhl-1* would lead to overt structural alterations of the cuticle, we investigated different reporters of cuticle structure. However, we did not observe any apparent changes for MUP-4::GFP (transmembrane protein required for epithelial cell and cuticular matrix attachment), AJM-1::GFP (superficial shape of seam epidermis), and SCM::GFP (seam cells, responsible to synthesizing the alae, distinctive region above the seam cells) (**Suppl. Fig. 2D-F**). Taken together, these data suggest that *vhl-1(ok161)* related expression changes in collagen genes may - while lacking gross changes in cuticle morphology - result in functional cuticle permeability defects.

### Hypodermal expression of VHL-1 partially rescues body length

We went on to confirm that the phenotype observed was indeed mediated by loss of *vhl-1* and to address the question whether expression of *vhl-1* was required in specific cell types regarding the ECM phenotype. Using a transgene driven by the *vhl-1* promoter fully rescued body length in *vhl-1* mutants and overexpression of VHL-1::GFP did not affect morphology in wildtype worms (**Fig. 4A** and **4B**). VHL-1::GFP was expressed all over the body with a subcellular localization that was not restricted to the cytoplasm but also present in the nucleus (**Suppl. Fig 4A**). Ubiquitous overexpression of VHL-1::GFP in *vhl-1(ok161)* worms also lead to the recovery of the expression of genes highly regulated upon loss of *vhl-1* (**Fig. 4C**). None of the tissue-specific promoters (*elt-2p* – intestine; *unc-119p* – neurons; *dpy-7p* hypodermis) driving *vhl-1*::GFP expression could recover the size of the animals (**Suppl. Fig 3B-D**); although a trend was seen for hypodermis-specific expression. However, a full rescue indeed depended on more widespread expression as driven by the endogenous promoter of *vhl-1*. Therefore, we can conclude that *vhl-1(ok161)* mutants’ abnormal size is due to *vhl-1* loss of function, and that no singular tissue-specific rescue is able to restore body size, which is only achieved with full body expression.

**Figure 4:**
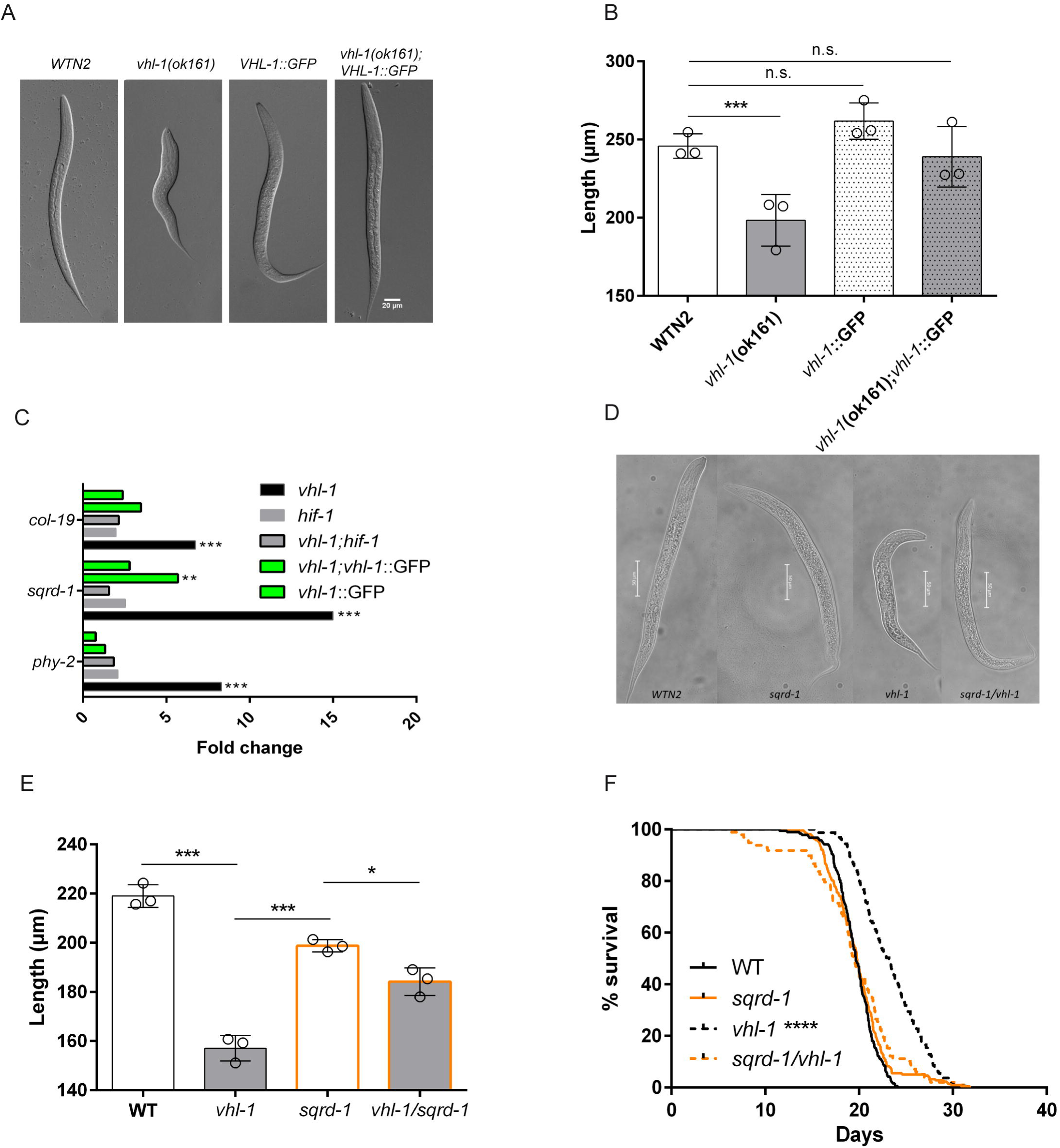
*sqrd-1* is partially responsible for *vhl-1(ok161)* phenotypes. **A)** Representative images of L1 *vhl-1(ok161)* rescue by *vhl-1*p::*vhl-1* construct; **B)** Size quantification of *vhl-1(ok161)* L1 mutants and *vhl-1p*::*vhl-1*::GFP rescue; **C)** mRNA expression levels of three strongly upregulated genes in *vhl-1(ok161)* in comparison to *hif-1(ia4)* and *vhl-1p*::*vhl-1*::GFP rescue; **D)** Size quantification of F1 generation L1 worms after *hif-1* RNAi exposure in P0; **E)** Size quantification of different L1 mutant animals, including *sqrd-1* (mr28) and double mutants *vhl-1(ok161)*/*sqrd-1* (mr28); F) Lifespan of different mutants. N = WT: 198, *vhl-1*: 278, *sqrd-1:* 203, *vhl-1/sqrd-1*: 317 Significance was calculated by comparing the mean value of the three independent experiments by a One-Way ANOVA followed by a Tukey post-hoc or a survival Kaplan-Meier analysis. * = p<0.05, ** = p<0.01, *** = p<0.001, **** = p<0.0001, ns = non-significant

### *sqrd-1* contributes to the effect of HIF-activation on body length

Next, we aimed to pin down downstream effectors underlying the impact of *vhl-1* mutation on morphology. Based on the RNAseq data, we selected 20 genes strongly positively regulated in *vhl-1* mutants in a *hif-1* dependent manner. Based on RNAi clone availability, these genes were included in an RNAi screen for body size (**Suppl. Table 1**). To check if the phenotype was reversible at all by RNAi, we first performed an experiment using *hif-1* RNAi. Notably, this approach (in contrast to the mutant) only partially rescued the length of the animals (**Suppl. Fig. 3E**). Therefore, partial rescue of size was considered as the benchmark for our screen, and we proceeded with 15 candidate genes. The knockdown of one of these genes, partially, but consistently (n=3), recovered length, the Sulfide Quinone Oxireductase (SQOR) *sqrd-1* (**Suppl. Fig. 3E**).

The recovery of the short phenotype of *vhl-1(ok161)* worms was considerably higher when we crossed them with a loss of function mutant of *sqrd-1* (mr28) (*vhl-1* 155µm x 186µm *vhl-1/sqrd-1*) (**Fig. 4D** and **E**). *sqrd-1* (mr28) worms do not differ from WT regarding body length. *sqrd-1* was indeed also one of the most strongly regulated genes in our RNAseq dataset (*vhl-1* vs WT log2 Fold Change: 3,64909, p=0,00005). Its expression levels, together with 2 ECM-associated genes chosen from the RNAseq dataset were fully restored by the overexpression of *vhl-1* in the *vhl-1*p::*vhl-1*::GFP transgenic animal (**Fig. 4C**). Taken together our data suggest that *sqrd-1* is a *hif-1* target that is at least partially responsible for the body morphology phenotype of *vhl-1(ok161)* mutant worms. Collagen had previously been linked to longevity in the nematode model (20) and *vhl-1* mutant worms are known to be long-lived (17). Interestingly, when measuring lifespan, we observed that the long lifespan of *vhl-1(ok161)* animals was completely reversed back to WT in the *vhl-1/sqrd-1* double mutant (**Fig. 4F**). Taken together these data suggest that *sqrd-1* is be an important downstream effector of *vhl-1* KO not only regarding body size, but also lifespan.

### The effect of sqrd-1 regarding body length and longevity of vhl-1 mutant worms depends on col-88

Considering that *sqrd-1* may mediate its effects on body morphology through collagens, we selected the top 5 most upregulated collagen genes in our RNAseq data of *vhl-1(ok161)* worms (**Suppl. Table 2**), and analyzed them for body size and *sqrd-1* epistasis by the means of mRNA expression. While knockdown of three of the collagen genes were able to slightly, but significantly increase *vhl-1* worm size (**Fig. 5A**), only one of these was differentially regulated in a manner that suggested epistasis with *sqrd-1* and *vhl-1,* being upregulated in the *vhl-1(ok161),* and downregulated in *sqrd-1* (mr28) and *vhl-1/sqrd-1* double mutant (**Fig. 5B**): *col-88* (**Fig. 5A** and **B, Suppl. Fig. 4A**). Indeed, RNAi-mediated knockdown of *col-88*, resulted in an indistinguishable body length between *vhl-1* mutants and *vhl-1;sqrd-1* double mutant worms (**Fig. 5C**) and mean lifespan (**Fig. 5D**). In conclusion, *col-88* seems to be positively regulated by *sqrd-1*, which in turn is upregulated upon *vhl-1* deletion/HIF-1 activation, seemingly leading to reduced body size in *vhl-1*(ok161) mutants.

**Figure 5:**
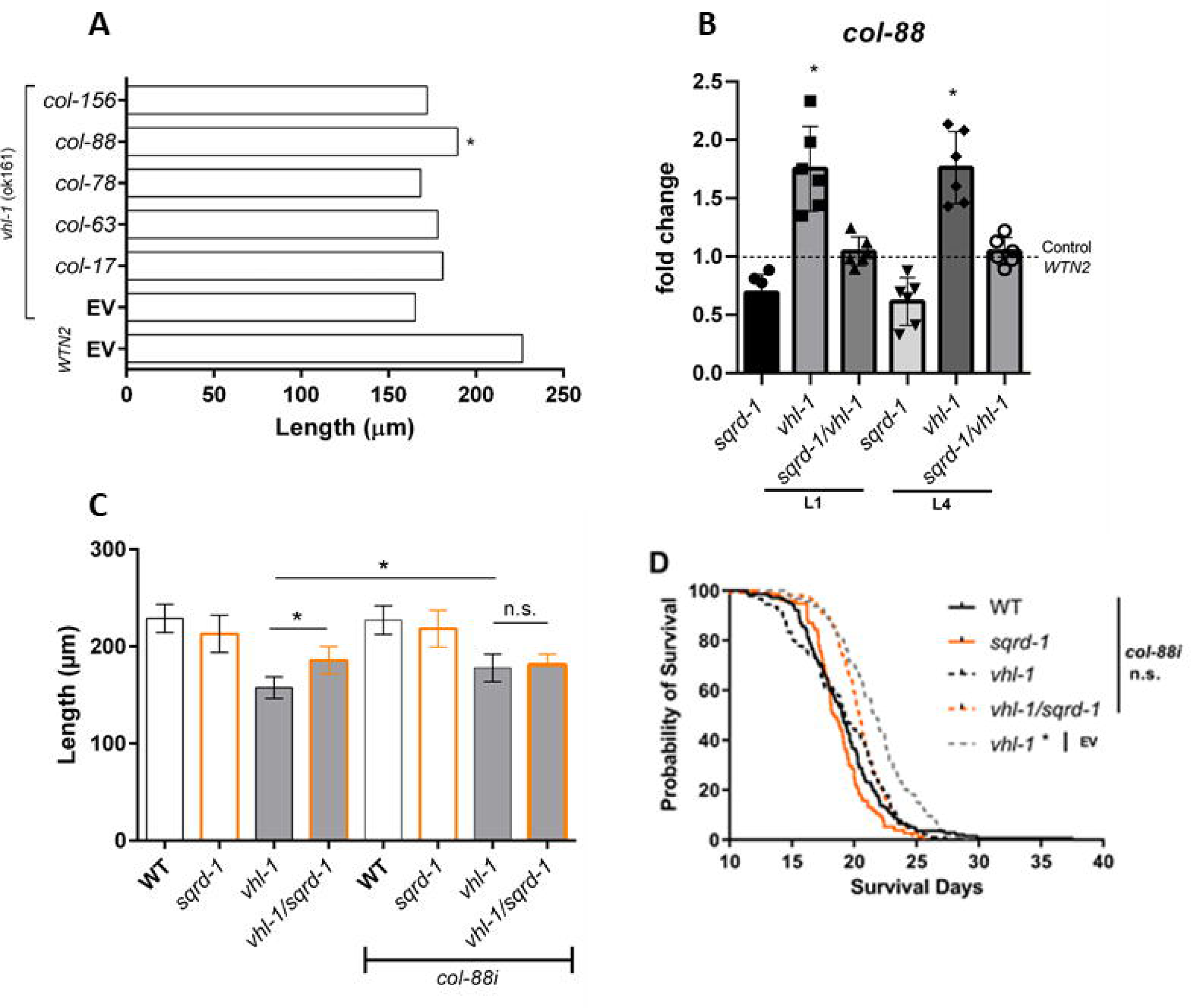
*col-88* acts downstream of *sqrd-1* regarding body length and longevity of *vhl-1* mutant worms. **A)** Body length upon exposure of P0 to 5 RNAi collagen clones that were upregulated in *vhl-1(ok161)*, followed by wormsizer analysis on F1 L1 newly hatched worms**; B)** mRNA expression levels of *col-88* in L1 and L4 animals in different genetic backgrounds; **C)** Body length of F1 generation L1 mutant worms after *col-88* RNAi exposure in P0; **D)** Lifespan of different mutants in RNAi bacteria for *col-88* or empty vector (L4440). N= WT 245, *sqrd-1* 308*, vhl-1* 289, *vhl-1/sqrd-1* 277, *vhl-1* (EV) 146. Significance was calculated by comparing the mean value of the three independent experiments by a One-Way ANOVA followed by a Tukey post-hoc. * = p<0.05, ** = p<0.01, *** = p<0.001, ns = non-significant

## Discussion

Here we present evidence for the link in between the hypoxia pathway regulator von Hippel-Lindau, VHL/*vhl-1,* and collagens to ultimately control body size and lifespan. Our data points to the participation of a sulfide metabolism related protein, the sulfide quinone oxidoreductase 1 SQOR/*sqrd-1,* as an important mediator to control these phenotypes. While sqrd-1 expression is induced by HIF-1, a specific collagen (*col-88*) appears to act downstream of *sqrd-1*.

pVHL has previously been implicated as a player in the regulation of ECM biology and integrity (11). However, a direct functional link has not been fully elucidated. Previous results pointed towards ECM association being a mainly HIF-independent phenotype. In fact, VHL has been shown to act independently of HIF-1 when it comes to its interaction with collagen 4 alpha 2 (COL4A2) and its proper assembly into the extracellular matrix (27). In contrast, our data clearly show HIF-dependence for both body length and ECM integrity. However, our results do not rule out HIF-independent ECM-associated roles of pVHL. Interestingly, *vhl-1;hif-1* double mutants appeared to be slightly shorter than WT or hif-1 mutant worms. Yet, this difference did not reach statistical significance anymore, so future studies will need to examine this question in more detail The abnormal Hoechst uptake – as well as the paraquat sensitivity – point towards defects in cuticle integrity, a finding that has been previously observed upon deletion of collagens *dpy-9* and *dpy-10* (28). However, our results cannot rule out that intestinal leakiness contributes to this result. Again, such a phenotype may involve changes in collagen composition of the basement membrane, but may also be a consequence of defective intestinal cell-cell junctions (29,30)).

The small body size persists through several rounds of molting and is independent of starvation, since larval stage 4 animals (not starved) display a short phenotype which is recovered upon *hif-1* (ia) deletion (**Fig. 1C**). Not only regular larval stages, but the radically different quiescent dauer larvae size is also changed by *vhl-1(ok161)* mutation. This finding indicated that the causal factor for body changes had to be a widely common biomolecule, such as structural parts of the cuticle, such as collagens. In that direction, *cut-6* knockdown, predicted to encode a structural part of the cuticle, induces similar changes in dauer larvae, resulting in a dumpy phenotype (31). In this regard, collagens are not only a major component of *C. elegans* cuticle (25), but are associated with well-studied worm phenotypes associated with short body length, such as Sma (small) and Dpy (dumpy) (25,32), and Dpy associated genes were in fact regulated in *vhl-1(ok161)* mutants in a *hif-1(ia4)* dependent manner (**Suppl. Fig. 1A**). The induction of collagens upon loss of *vhl-1* could be recapitulated on the protein level, both when quantifying overall collagen levels or *col-19* as an example for one specific collagen. Importance of these changes for ECM biology is not only shown by the change in body length but also reflected by defects in cuticle integrity in *vhl-1* mutant worms.

Using an RNAi screen for downstream mediators of the body length reduction among HIF-induced genes (**Suppl. Table 1** and **Suppl. Fig. 3E**), we identified a rather unexpected gene, the sulfide quinone oxireductase 1, *sqrd-1*. *sqrd-1* is the orthologue of mammalian SQOR, and encodes a mitochondrial sulfide quinone oxidoreductase responsible for the first step of the conversion of sulfides into less toxic persulfides (33,34). In *C. elegans* it has been described to be required for survival under toxic concentrations of hydrogen sulfide (H_2_S) (34), meaning its deletion hinders *C. elegans* sensitive to this gas, similarly to HIF-1 deletion (35,36). Interestingly, one previous study found that HIF-1 activation after H_2_S exposure was exclusive to the hypodermis and independent of *vhl-1* (35).

Even though link has not been fully understood, the relationship between H_2_S and ECM remodeling has been documented. As an example, spontaneously hypertensive rats treated with H_2_S donor Na_2_HS did not produce as much collagen 1, collagen 4, and hydroxyproline in their vascular smooth muscle cells (37). Similarly, H_2_S deficiency is key to ECM remodeling in diabetic kidney, whereas treatment with the H_2_S donor GYY4137 is able to repress collagen 1 and 4 overexpression in a mouse model (38). Additionally, pulmonary and liver fibrosis progression seem to also have endogenous H_2_S levels decreased, whereas the treatment with donors can have mild ameliorative effects (39,40). Interestingly, hypoxia itself, as well as pVHL inactivation in ccRCC, are known to correlate with buildup of intracellular H_2_S (41–44)H_2_S is considered a modulator of HIF-1 (45–47), and may contribute to tumor survival and progression (41). Whether this gasotransmitter is a consequence or just a collateral in these scenarios is yet to be clarified, since the regulation of the pathway that controls the production of this gas, and even small molecules that are able to produce it at variable rates, amounts, and with cellular compartment specificity, could be further explored as potential therapeutic targets for diseases that involve HIF-1 activation, hypoxia, or pVHL inactivation (38,48). In that sense, we believe that our work sheds further light onto this topic, demonstrating the necessity of a sulfide catabolism related gene in an ECM-related phenotype regulated by HIF-1.

*C. elegans* possesses 181 collagen-coding genes(49), and the redundancy in between them is not as big as one would expect. A recent study screened 93 collagen genes by means of RNAi and only 6 of these genes were associated to a cuticle barrier defect (28). Knockdown of two of these genes, *dpy-9* and *dpy-10,* did not only made the cuticle permeable, but also increased the uptake and toxicity of the stressor paraquat over adult animals. Consequently, increased stress resistance may also be one of the links between collagens and lifespan extension.

*Vhl-1(ok161)* mutant worms are known to be both stress resistant and long-lived (17), and longevity in this strain is mediated by HIF-1. Interestingly, an unexpected link between collagens and longevity had previously been established in long-lived *daf-2* (e1370) mutants (20). This mechanism is not specific to longevity mediated by inhibition of insulin signaling but was also shown for caloric restriction induced by *eat-2* mutation, or germline ablation via *glp-1* knockout (20). Interestingly, one of the collagens examined in our study, *col-19*, was not induced before later adulthood. In line with the direct connection between collagen expression and lifespan, we could now show that *sqrd-1* dependent induction of a specific collagen – col-88 – was required for the lifespan extension in *vhl-1* mutant worms.

Based on its amino acid sequence COL-88 is predicted to be structural part of the cuticle, and it was first identified in a RNAi screen for osmotic hypersensitivity (50). To the extent of our knowledge, no other study has described any specific COL-88 function or associated phenotype. We believe that the upregulation of this specific collagen plays an important role in *vhl-1* body size, probably playing a role in ECM overall organization. SQRD-1 is central to the metabolism of H_2_S. Since H_2_S is an important mediator of lifespan in different species under caloric restriction (51), and H_2_S donors are known for improving stress resistance and lifespan in *C. elegans* (52–54), one could speculate that the effects of this gasotransmitter on lifespan and stress resistance is also ECM remodeling, consequently affecting body size. Recently, it has been shown that H_2_S donors targeted specifically to the mitochondria can induce collagens (Type I) on the protein and mRNA level in normal human dermal fibroblasts (NHDFs) and mouse skin in vivo to protect against UV induced damage in an Nrf2 dependent manner (55).

Novel elasticity measurements have recently revealed that older worms become ‘’stiffer’’ while long lived *daf-2* animals were still more elastic in their cuticle structure at the same age, while the knockdown of *col-120* was able to ablate *daf-2* elasticity and lifespan (20,56). Similarly, collagens seem to be the most significantly altered group of proteins in the proteome of the long-lived germline loss model *glp-1* (e2141) (57), where, interestingly, H_2_S levels are increased, and its main source of production, the transsulfuration pathway, is required for lifespan extension (58). Taken together, our data and the studies mentioned point to a possible new link in between H_2_S metabolism, ECM composition and longevity. It will be intriguing to further characterize this link in future studies and to extend the work to mammalian models. Such knowledge will be of special interest regarding the role of the pVHL/HIF-axis in tumorigenesis considering the importance of the ECM to tumor growth and metastasis.

## Materials and Methods

### *C. elegans* strains and maintenance

The following strains have been used in this study: *C. elegans* Bristol N2 (WT); CB5602 [*vhl-1(ok161)* X]; ZG31 [hif-1(ia4)]; AR13 [*sqrd-1* (mr28) IV]; TMB031 [*hif-1*(ia4) V; *vhl-1*(ok161) X]; CB1370 [daf-2 (e1370) III]; TP12 [kaIs12 (*col-19*::*gfp*)]; JR667 [*unc-119 (e2498::Tc1) III; wIs51* (*SCMp::*gfp*;unc-119+)*] V]; TBM033 [*vhl-1(ok161)* X; kaIs12 (*col-19*::*gfp*]; TMB036 [*jcIs* (*ajm-1*::*gfp*;*unc-29*+;*rol-6* (su1006) IV; *vhl-1(ok161)* X]; AA813 [*jcIs1* (*ajm-1*::*gfp*;*unc-29*+;*rol-6* (*su1006*) IV]; TMB101 [*vhl-1(ok161)* X; *sqrd-1* (mr28) IV]. All strains were provided by the *Caenorhabditis* Genetics Center (University of Minnesota, Twin Cities, MN, USA). CGC strains have been outcrossed into our WT background at least 2x. All strains named under “TMB” have been generated in our lab. Genotypes were confirmed by using the primers included in **Suppl. Table 3**.

Worms were handled and maintained at 20 °C on NGM (nematode growth medium)/*Escherichia coli* OP50 Petri dishes. NMG was prepared with 3g/L of NaCl, 2,5g/L of Bacto Peptone, and 18g/L of Bacto Agar. After autoclaving 25mM of KPO_4_, 0.005mg/mL Cholesterol, 1mM MgSO_4_, and 1mM of CaCl_2_ were added. *E. coli* OP50 bacteria were grown overnight (12-16h) from a single colony. O.D. was adjusted to 1.5 with LB media (20g/L) by measuring density at 650nm.

Age synchronized populations used for experiments were obtained by isolating embryos from gravid hermaphrodites using bleaching solution (1% NaOCl; 0.25 M NaOH), followed by neutralization with M9 buffer (KH_2_PO_4_ 12g/L, Na_2_HPO_4_ 24g/L, NaCl 20g/L, MgSO_4_ 1mM). The axenization of the worms has been done by employing the same bleaching solution over a small group of pregnant hermaphrodites if contamination was spotted. All experiments were conducted at 20±0.5°C in a humidified-controlled environment. No ethical approval was needed.

### RNAi

*E. coli* HT115 (DE3) transformed with the L4440 vector, empty or including dsRNA specific to the gene which knockdown is intended, were inoculated in LB medium containing 100μg/mL of ampicillin and 12,5μg/mL of tetracycline overnight. The next day, the medium was concentrated to a 600nm O.D. of 1,5, upon which 1mM of the expression inducer IPTG (isopropyl β-D-1-thiogalactopiranoside) was added, besides being already present at the same concentration inside the NGM medium in which bacteria were seeded as a food source for *C. elegans*.

### Body size measurement

Age synchronized L1 populations were obtained by bleaching a large plate with pregnant hermaphrodites (D0) either on regular OP50 *E. coli* (experiments comparing different knockouts) or fed with HT115 (DE3) *E. coli* carrying different RNAi clones (for experiments carried after knockdown). These populations have either been allowed to grow until they reached the L4 (48h after bleaching) stage or placed in unseeded plates so that L1 animals were measured (16h after bleaching). The WormSizer plugin was used to analyze the length and width of the worms. Worms were imaged at 63x magnification using an Axiozoom (Zeiss) microscope. At least 50 worms per group were imaged per biological replicate. Measurements were manually reviewed after WormSizer analysis to exclude potential confounders (e.g. any debris, bacterial lawn, or incorrectly selected worm).

### Dauer formation assay

Age synchronized 500 L1 animals were placed either in NGM plates containing OP50 or no food source. OP50-containing plates were kept at 20°C and plates with no food source were kept at 25°C to induce dauer formation. After 3 days we have washed the animals with M9 buffer and treated them with 1% SDS for 30min. After the treatment the animals were washed 3x with M9 buffer and placed in a OP50-containing NGM plate and let to rest for 24h. The animals were scored and the percentage of survival was estimated based on the initial number of animals placed in each plate (500).

### Preparation of RNA samples, RNA extraction and sequencing

Synchronized worms were grown according to their optimum growth requirements and collected at their D1 adult stage in Trizol (Invitrogen) in three independent biological repeats. Total RNA was extracted using RNeasy Mini spin column (QIAGEN). Concentration and purity of the RNA was measured by NanoDrop. polyA+ mRNA was isolated from total RNA with NEBNext Poly(A) mRNA Magnetics Isolation Module (New England Biolabs). RNA-Seq libraries were prepared with NEBNext Ultra Directional RNA Library Prep Kit for Illumina (New England Biolabs). Capillary electrophoresis (Agilent Bioanalyser and Agilent Tapestation) was used to assess quality and quantity at all steps. Libraries were quantified by fluorimeter, immobilized and processed onto a flow cell with a cBot (Illumina) followed by sequencing-by-synthesis with TruSeq v3 chemistry on a HiSeq2500 performed at the Max Planck Genome Center CGC (Cologne, Germany).

### Bulk RNA sequencing analysis

The reads were trimmed with Trimmomatic (version 0.36) (59) using default parameters. The trimmed reads were mapped to the WBcel235 Caenorhabditis elegans reference genome with STAR version 2.6.0 (60) using default parameters. The analysis of differential gene expression was performed using the R package DeSeq2 (version 1.26.0) (61,62). For visualization purposes the batch effect was removed using the method removeBatchEffect from the R package limma (63). The enrichment analysis was done using the R package topGO (64,65). For the heatmap and clustering the R package pheatmap was used (66).

Bulk RNA sequencing data are available in the ArrayExpress database (https://www.ebi.ac.uk/biostudies/arrayexpress) under accession number E-MTAB-13659.

### Collagen analysis

Synchronized L1 larvae were placed on 9 cm NGM plates containing OP50 bacteria at 20°C and kept there till they reached the L4 stage (48h). Larvae were washed from the plates and transferred to new 5-Fluoro-2’deoxyuridine (FUdR) containing plates till day 1 or day 8 of adulthood. At those time points, the remaining animals were harvested by washing 3x with M9 buffer. 3000 worms per strain and condition were used for the assay. Collagen levels were determined using the *QuickZyme* Biosciences Total Collagen Kit (QZBTOTCOL1), which detects Hydroxyproline, according to the manufacturer’s instructions. Briefly, hydroxyproline is quantified at 570nm after being conjugated with chromophore. Values are displayed as μg/mL.

### Western blotting

Western blot analysis was used to measure the expression of the GFP reporter. First, 50 worms were picked into 1 ml lysis buffer and sonicated ten times for 30 seconds. Afterwards 2xLaemmli buffer (150 mM TrisHCl pH 6,8, 4% SDS, 20% Glycerol, 0.002% Bromophenol blue, 0.1 M DTT) was added and the sample was incubated for 5 minutes at 95°C. 30 µl of the samples were loaded on a 4 to 12% Bis-Tris polyacrylamid gel and the gel was run at 200 V constant for 60 minutes. Gels were transferred to PVDF membranes. Staining was performed using an anti GFP antibody (Santa Cruz, sc-8334, 1:5,000) and an anti-rabbit secondary antibody (1:15,000, Jackson ImmunoResearch, 711-165-152). Visualization was performed using Femto Luminol (Invitrogen) and a chemiluminescence imaging system (FusionSolo, PeqLab).

### Epifluorescence visualization and image acquisition

A Zeiss Axiovert 200 microscope (Zeiss, Jena, Germany) equipped with a Plan Apochromat 20x objective and a GFP filter set was used for fluorescence microscopy. All pictures were taken using this microscope, except for worms treated with the Hoechst staining for which the confocal microscope (Zeiss Meta 710, Type Confocal Laser Scanning Microscope 20x objective) was used. The AXIO Zoom.V16 was used to take the pictures for the WormSizer plug-in and to pick GFP fluorescent worms. The AXIO Zoom.V16 was equipped with an objective from ZEISS (PlanNeoFluar Z 1x/0.25 FWD 56 mm), with a filter set at 74 HE, and with a Leica DFC450 camera. Worms were mounted onto 2% agar pads and paralyzed with 15mM levamisole. Fluorescence measurement was calculated as CTCF (Corrected Total Cell Fluorescence).

### Cuticle permeability assay

To analyze the permeability of the worms’ extracellular matrix, Hoechst staining was performed as previously described (67). Briefly, young adults were placed in 1.5 ml tubes with 1 ml M9 buffer containing 1μg/ml Hoechst 33342 and incubated for 15 minutes at RT. After washing three times in M9, worms were immobilized using 60 mM Sodium Azide on 2% agarose pads and imaged at the confocal microscope keeping acquisition parameters constant for intensity comparison. The tail region was analyzed to access oral/intestinal absorption, while the head region was analyzed to access cuticle absorption.

### Generation of transgenic lines by DNA bombardment

Transgenic lines for reexpression of *vhl-1* under its endogenous promoter were generated by bombarding *unc-119 (ed3)* worms with plasmid harboring the *vhl-1*::GFP transgene under the control of *vhl-1* promoter and *unc-119* as a selection marker. 30mg of gold particles (Gold powder 0.3-3micron/99.9% from chemPUR) were transferred in a 1.5mL tubes (DNA LoBind, Eppendorf), vortexed with 1mL 70% EtOH for 5min, incubated for 15min at room temperature, and shortly spun down so that the supernatant could be discarded. Gold particles were washed three times with 1mL of water and resuspended in 500µL of 50% glycerol for a 60mg/mL concentration of the particles. This stock solution was coated with plasmid DNA containing the construct of interest by vortexing it for 5min and mixing 20µL of its supernatant with 10µL of 1-2mg/mL solution of plasmid DNA and vortexing the mix for 1min. 300µL of 2.5M CaCl_2_ was added over the mix, vortexed for 1min, followed by 300µL of 2.5M CaCl_2_ and 3 more minutes of vortexing. The particles were centrifuged at 12000rpm for 1min and washed with 800µL of 70% EtOH, centrifuged again and washed with 500µL of 100% EtOH, and finally washed one last time after centrifugation with 80µL of 100% EtOH. This solution was distributed on 7 macrocarriers and allowed to dry.

Briefly, a mixed population of adults (D0-D2 adults) were cultured in EP (peptone enriched) plates, collected using M9 and further transferred to cold unseeded 10cm NGM plates. Plates with animals were placed in the gene gun (Model PDS-1000/He Biolistic Particle Delivery from BIO-RAD) with gold particles that were previously coated with *vhl-1p::vhl-1::*GFP plasmid DNA. After bombardment the worms were washed off the plates and distributed on 10cm NGM plates seeded with OP50 *E. coli* and incubated at 20°C: After two weeks only the worms containing the *unc-119* rescue, and consequently our construct of interest, were able to enter the dauer stage due to starvation. We confirmed transgene presence by detecting GFP by microscopy and PCR.

### Plasmid construction and microinjection

Tissue specific rescue was achieved by cloning chosen promoters into pdest-MB14 plasmid, which includes GFP in frame. Briefly, *vhl-1* entire genomic sequence was cloned plus a 50bp region upstream of the *vhl-1* first codon, which should cover its promoter. All primers used are included in **Suppl. Table 3**.

The injection was performed as described before (68) using an Axio Observer A1 (Zeiss [Carl Zeiss], Tornwood, NY) and FemtoJet (Eppendorf, Hamburg, Germany) microinjection system. Briefly, five gravid adult worms were placed on a coverslip with a dried 2% agarose pad on it and were covered with a layer of Halocarbon oil (Sigma, St Louis, MO) to prevent drying during the process. The injection needle (Femtotips II, Eppendorf) was filled with a mix of 5µg of the plasmid of interest and 1µg of mCherry as an injection control, all diluted with Water for Injection (Ampuwa). Each worm was injected in one of the two gonads, and at least 30 worms were injected with each specific plasmid. After injection the worms were immediately placed on a NGM plate seeded with OP50 to recover, and incubated at 20°C.

### Quantitative Real Time PCR

Briefly, RNA extraction was performed with the Direct-zol RNA Miniprep kit (Zymo Research) following the manufacturer’s instructions. Prior to the reverse transcription by using the High-Capacity cDNA Reverse Transcription kit (Applied Biosystems), RNA concentration and sample quality were assessed on a Nanodrop spectrophotometer (Peqlab). mRNA was assessed by SYBR Green (ThermoFisher Scientific) qPCR using *act-1* as endogenous control. Primers are listed in Supplementary Table S3. The qPCR experiments were performed on a QuantStudio 12LK Flex Real-time PCR System (ThermoFisher Scientific). All primers used are included in **Suppl. Table 3**. For data analysis, all results were normalized to the housekeeping gene *act-1* using the delta-delta CT followed by a One-way ANOVA and a Tukey posthoc test (*p*L<L0.05).

### Lifespan analysis

All lifespan analyses were conducted using the Lifespan Machine (69) For each lifespan experiment 35 young adult worms were transferred to 60 mm NGM or RNAi plates containing 100µm Fluorodeoxyuridine (FUdR) to inhibit reproduction, or additionally Paraquat (Methylviologen, Sigma) 10mM for stress resistance. For each strain, at least 5 plates were used. The plates were put upside down without the lids on the glass plate of the scanners and fit into a rubber cast (rubber mask) to prevent desiccation. The animals were scanned each 30 minutes until the end of the experiment (determined by the expected time at which all animals would be dead e.g. 30 days for *WT*). After the end of the experiment, a storyboard containing a timeline for every animal in the experiment was generated, so that their movement cessation could be manually checked, as well as any other discrepancy (e.g. animal confused with artifact). The final death events of the worms were collected in a table and plotted as a survival curve.

### Statistical analysis

The software GraphPad Prism 9 was used for all statistical analysis conducted. *, **, ***, and **** represent p values that were lower than 0.05, 0.01, 0.001, and 0.0001 respectively. The specific statistical tests used are specified for each experiment in the figure legends and in the respective methods section.

## Acknowledgments

We thank Serena Greco-Torres for excellent technical support and WormBase for curated gene and phenotype information (https://wormbase.org/). Some strains were provided by the *Caernohabditis elegans* Genetics Center (CGC), which is funded by NIH Office of Research Infrastructure Programs (P40 OD010440). This study was supported by the German Research Foundation (DFG MU 3629/6-1 to R.-U.M.). C.Y.E. and C.S. received funding from the Swiss National Science Foundation Funding (SNF P3 Project 190072).

## Author Contributions

All authors participated in analyzing and interpreting the data and reviewed the manuscript. R.E., R.-U.M, and W.S. designed the experiments. R.E., M.N. and W.S. performed experiments. K.B., A.B. and M.C. analyzed the RNAseq data. W.S. wrote and R.E., M.I. and R.-U.M. reviewed the original manuscript. C.Y.E., S.B. and C.S. supported the setup of the lifespan machine and C.Y.E. provided scientific advice regarding the Collagen experiments R.-U.M, M.I. and F.F. supervised the work supervised the project and R.-U.M. provided funding.

## Supporting information captions

**Suppl. Figure 1: Regulation of genes associated with common ECM related phenotypes**

Expression change of genes associated with the dumpy phenotype compared to WT; **B)** Respective analysis for genes associated with the long phenotype; **C)** Respective analysis for genes associated with the roller phenotype; **D)** Number of dauer animals measured by % of survival after treatment with SDS after normal culture (20°C) or dauer inducing culture (starving at 25°C). The group of genes associated with each phenotype was taken from the Matrisome project (22). Boxes and whiskers represent the 25-75th percentile and the minimum to maximum values, respectively. Significance was calculated by comparing the mean value each group with a One-Way ANOVA followed by a Tukey post-hoc. * indicates a p<0.05 for the comparison in between the signaled groups.

**Suppl. Figure 2: Collagen reporter expression, stress resistance, and cuticle associated reporters**

Representative images of the blotting against GFP in *col-19*::GFP WT and *vhl-1(ok161)* backgrounds; **B)** Densiometry measurements of A**; C)** Survival curve of animals exposed to paraquat 10mM**; D)** Representative pictures of the MUP-4::GFP reporter**; E)** Representative pictures of the AJM-1::GFP reporter; **F)** Representative pictures of the SCM::GFP reporter. Significance was calculated by comparing the mean value each group with a T-student test or a survival Keplan-Meier analysis. *** indicates a p<0.001 for the comparison in between the signaled groups.

**Suppl. Figure 3: Tissue-specific rescue of *vhl-1*::GFP expression and RNAi screen for size regulators**

**A)** Representative pictures of L1 worms expressing *pvhl-1::vhl-1*::GFP; **B)** Size of *vhl-1(ok161)* worms expressing *vhl-1::*GFP in the intestine driven by the *elt-2* promoter; **C)** Size of *vhl-1(ok161)* worms expressing *vhl-1::*GFP in the neurons driven by the unc-*119* promoter; **D))** Size of *vhl-1(ok161)* worms expressing *vhl-1::*GFP in the cuticle driven by the *dpy-7* promoter; **E)** Size of *vhl-1(ok161)* F1 worms of which P0 was exposed to RNAi against the indicated genes, including *sqrd-1* in red. Significance was calculated by comparing the mean value each group with a One-Way ANOVA followed by a Tukey post-hoc or t-test Student when only two groups were present. * indicates a p<0.05 for the comparison of any group with vhl-1(ok161) alone or in empty vector (EV) RNAi.

**Suppl. Figure 4: Expression of different collagen candidates**

Expression of *col-17, col-63, col-78*, and *col-88, and col-156* in *vhl-1(ok161)*, *sqrd-1(ar13)*, and *sqrd-1/vhl-1* double mutant. mRNA amounts is expressed as fold change compared to the WT expression levels (dotted line) of each respective collagen. Significance was calculated by comparing the mean value each group with an One-Way ANOVA followed by a Tukey post-hoc.

**Suppl. Table 1: RNAseq data from *vhl-1(ok161)*, *hif-1(ia4)*, and *vhl-1;hif-1* double mutant.**

Values are expressed as fold change (log2), whereas green values signal significant upregulation, red signals significant downregulation, and white values signal no change compared to *WT* controls. Statistically significant differences between WT and the other groups was calculated the R package DeSeq2.

**Suppl. Table 2: RNAseq data regarding the top regulated collagen genes in *vhl-1(ok161)* mutants:**

Values are expressed as fold change (log2), whereas green values signal significant upregulation, red signals significant downregulation for the described comparison. Statistically significant differences between WT and the other groups was calculated calculated the R package DeSeq2.

## References

1. Semenza GL. Hypoxia-Inducible Factor 1 (HIF-1) Pathway. Science’s STKE. 2007 Oct 9;2007(407).

2. Maxwell PH, Wiesener MS, Chang GW, Clifford SC, Vaux EC, Cockman ME, et al. The tumour suppressor protein VHL targets hypoxia-inducible factors for oxygen-dependent proteolysis. Nature [Internet]. 1999;399(6733):271–5. Available from: 10.1038/20459

3. Ohh M, Park C, Ivan M, Hoffman M, … TKN cell, 2000 undefined. Ubiquitination of hypoxia-inducible factor requires direct binding to the β-domain of the von Hippel–Lindau protein. nature.com [Internet]. [cited 2023 May 2]; Available from: https://idp.nature.com/authorize/casa?redirect_uri=https://www.nature.com/articles/ncb0700_423&casa_token=689Z3WxIJroAAAAA:ba2RFWrpRhZyuvawX9HllUy4cakzU7fODNqRQEkiA869oiQsVUCTt19RW59fyotW4piBIwIA9V7hKGALxg

4. Maxwell PH, Wlesener MS, Chang GW, Clifford SC, Vaux EC, Cockman ME, et al. The tumour suppressor protein VHL targets hypoxia-inducible factors for oxygen-dependent proteolysis. Nature 1999 399:6733 [Internet]. 1999 May 20 [cited 2023 May 2];399(6733):271–5. Available from: https://www.nature.com/articles/20459

5. Gnarra JR, Zhou S, Merrill MJ, Wagner JR, Krumm A, Papavassiliou E, et al. Post-transcriptional regulation of vascular endothelial growth factor mRNA by the product of the VHL tumor suppressor gene. Proc Natl Acad Sci U S A [Internet]. 1996 Oct 1 [cited 2023 May 2];93(20):10589–94. Available from: https://pubmed.ncbi.nlm.nih.gov/8855222/

6. Kaelin WG. Molecular basis of the VHL hereditary cancer syndrome. Nature Reviews Cancer 2002 2:9 [Internet]. 2002 Sep [cited 2023 May 2];2(9):673–82. Available from: https://www.nature.com/articles/nrc885

7. Reversion of deregulated expression of vascular endothelial growth factor in human renal carcinoma cells by von Hippel-Lindau tumor suppressor protein - PubMed [Internet]. [cited 2023 May 2]. Available from: https://pubmed.ncbi.nlm.nih.gov/8625303/

8. Na X, Wu G, Ryan CK, Schoen SR, Di’Santagnese PA, Messing EM. Overproduction of vascular endothelial growth factor related to von Hippel-Lindau tumor suppressor gene mutations and hypoxia-inducible factor-1α expression in renal cell carcinomas. Journal of Urology. 2003 Aug 1;170(2 I):588–92.

9. Ohh M, Yauch RL, Lonergan KM, Whaley JM, Stemmer-Rachamimov AO, Louis DN, et al. The von Hippel-Lindau tumor suppressor protein is required for proper assembly of an extracellular fibronectin matrix. Mol Cell. 1998;1(7):959–68.

10. Singh P, Carraher C, Schwarzbauer JE. Assembly of fibronectin extracellular matrix. Annu Rev Cell Dev Biol. 2010 Nov 10;26:397–419.

11. Stickle NH, Chung J, Klco JM, Hill RP, Kaelin WG, Ohh M. pVHL Modification by NEDD8 Is Required for Fibronectin Matrix Assembly and Suppression of Tumor Development. Mol Cell Biol. 2004 Apr 1;24(8):3251–61.

12. Russell RC, Ohh M. NEDD8 acts as a ‘molecular switch’ defining the functional selectivity of VHL. EMBO Rep. 2008 May;9(5):486–91.

13. Kurban G, Hudon V, Duplan E, Ohh M, Pause A. Characterization of a von Hippel Lindau Pathway Involved in Extracellular Matrix Remodeling, Cell Invasion, and Angiogenesis. Cancer Res [Internet]. 2006 [cited 2023 May 2];66(3):1313–22. Available from: www.aacrjournals.org

14. Kurban G, Duplan E, Ramlal N, Hudon V, Sado Y, Ninomiya Y, et al. Collagen matrix assembly is driven by the interaction of von Hippel-Lindau tumor suppressor protein with hydroxylated collagen IV alpha 2. Oncogene [Internet]. 2008 [cited 2023 May 2];27:1004–12. Available from: www.nature.com/onc

15. Vidović T, Ewald CY. Longevity-Promoting Pathways and Transcription Factors Respond to and Control Extracellular Matrix Dynamics During Aging and Disease. Frontiers in aging. 2022;3:935220.

16. Shen C, Shao Z, Powell-Coffman JA. The Caenorhabditis elegans rhy-1 gene inhibits HIF-1 hypoxia-inducible factor activity in a negative feedback loop that does not include vhl-1. Genetics [Internet]. 2006 [cited 2023 Apr 27];174(3):1205–14. Available from: https://pubmed.ncbi.nlm.nih.gov/16980385/

17. Müller RU, Fabretti F, Zank S, Burst V, Benzing T, Schermer B. The von Hippel Lindau tumor suppressor limits longevity. Journal of the American Society of Nephrology [Internet]. 2009 Dec 1 [cited 2023 Feb 23];20(12):2513–7. Available from: https://journals.lww.com/jasn/Fulltext/2009/12000/The_von_Hippel_Lindau_Tumor_Suppressor_Limits.9.aspx

18. Bishop T, Lau KW, Epstein ACR, Kim SK, Jiang M, O’Rourke D, et al. Genetic analysis of pathways regulated by the von Hippel-Lindau tumor suppressor in Caenorhabditis elegans. PLoS Biol [Internet]. 2004 Oct [cited 2023 Apr 26];2(10). Available from: https://pubmed.ncbi.nlm.nih.gov/15361934/

19. Statzer C, Jongsma E, Liu SX, Dakhovnik A, Wandrey F, Mozharovskyi P, et al. Youthful and age-related matreotypes predict drugs promoting longevity. Aging Cell. 2021 Sep;20(9):e13441.

20. Ewald CY, Landis JN, Abate JP, Murphy CT, Blackwell TK. Dauer-independent insulin/IGF-1-signalling implicates collagen remodelling in longevity. Nature [Internet]. 2015 Mar 3 [cited 2023 Feb 24];519(7541):97. Available from: /pmc/articles/PMC4352135/

21. Moore BT, Jordan JM, Baugh LR. WormSizer: High-throughput Analysis of Nematode Size and Shape. PLoS One [Internet]. 2013 Feb 22;8(2):e57142-. Available from: 10.1371/journal.pone.0057142

22. Teuscher AC, Jongsma E, Davis MN, Statzer C, Gebauer JM, Naba A, et al. The in-silico characterization of the Caenorhabditis elegans matrisome and proposal of a novel collagen classification. Matrix Biol Plus. 2019 Feb;1:100001.

23. Ewald CY. The Matrisome during Aging and Longevity: A Systems-Level Approach toward Defining Matreotypes Promoting Healthy Aging. Gerontology. 2020;66(3):266–74.

24. Teuscher AC, Statzer C, Pantasis S, Bordoli MR, Ewald CY. Assessing Collagen Deposition During Aging in Mammalian Tissue and in Caenorhabditis elegans. In 2019. p. 169–88.

25. Johnstone IL. Cuticle collagen genes. Trends in Genetics. 2000 Jan;16(1):21–7.

26. Kage-Nakadai E, Kobuna H, Kimura M, Gengyo-Ando K, Inoue T, Arai H, et al. Two Very Long Chain Fatty Acid Acyl-CoA Synthetase Genes, acs-20 and acs-22, Have Roles in the Cuticle Surface Barrier in Caenorhabditis elegans. PLoS One [Internet]. 2010 Jan 25 [cited 2023 Feb 24];5(1):e8857. Available from: https://journals.plos.org/plosone/article?id=10.1371/journal.pone.0008857

27. Kurban G, Duplan E, Ramlal N, Hudon V, Sado Y, Ninomiya Y, et al. Collagen matrix assembly is driven by the interaction of von Hippel–Lindau tumor suppressor protein with hydroxylated collagen IV alpha 2. Oncogene. 2008 Feb 7;27(7):1004–12.

28. Sandhu A, Badal D, Sheokand R, Tyagi S, Singh V. Specific collagens maintain the cuticle permeability barrier in Caenorhabditis elegans. Genetics [Internet]. 2021 Mar 31 [cited 2023 Feb 25];217(3). Available from: https://academic.oup.com/genetics/article/217/3/iyaa047/6123747

29. Zuo L, Kuo WT, Turner JR. Tight Junctions as Targets and Effectors of Mucosal Immune Homeostasis. Cell Mol Gastroenterol Hepatol. 2020;10(2):327–40.

30. Kim MR, Cho SY, Lee HJ, Kim JY, Nguyen UTT, Ha NM, et al. Schisandrin C improves leaky gut conditions in intestinal cell monolayer, organoid, and nematode models by increasing tight junction protein expression. Phytomedicine. 2022 Aug;103:154209.

31. Muriel JM, Brannan M, Taylor K, Johnstone IL, Lithgow GJ, Tuckwell D. M142.2 (cut-6), a novel Caenorhabditis elegans matrix gene important for dauer body shape. Dev Biol. 2003 Aug 15;260(2):339–51.

32. Myllyharju J. Collagens, modifying enzymes and their mutations in humans, flies and worms. Trends in Genetics. 2004 Jan;20(1):33–43.

33. Yong R, Searcy DG. Sulfide oxidation coupled to ATP synthesis in chicken liver mitochondria. Comparative Biochemistry and Physiology - B Biochemistry and Molecular Biology [Internet]. 2001 [cited 2023 Feb 26];129(1):129–37. Available from: https://pubmed.ncbi.nlm.nih.gov/11337256/

34. Marutani E, Morita M, Hirai S, Kai S, Grange RMH, Miyazaki Y, et al. Sulfide catabolism ameliorates hypoxic brain injury. Nat Commun [Internet]. 2021 Dec 1 [cited 2023 Feb 26];12(1). Available from: https://pubmed.ncbi.nlm.nih.gov/34035265/

35. Budde MW, Roth MB. Hydrogen sulfide increases hypoxia-inducible factor-1 activity independently of von Hippel-Lindau tumor suppressor-1 in C. elegans. Mol Biol Cell [Internet]. 2010 Jan 1 [cited 2023 Feb 26];21(1):212–7. Available from: https://www.molbiolcell.org/doi/10.1091/mbc.e09-03-0199

36. Miller DL, Budde MW, Roth MB. HIF-1 and SKN-1 coordinate the transcriptional response to hydrogen sulfide in Caenorhabditis elegans. PLoS One [Internet]. 2011 Sep 29 [cited 2023 Feb 26];6(9). Available from: https://pubmed.ncbi.nlm.nih.gov/21980473/

37. Zhao X, Zhang LK, Zhang CY, Zeng XJ, Yan H, Jin HF, et al. Regulatory effect of hydrogen sulfide on vascular collagen content in spontaneously hypertensive rats. Hypertens Res [Internet]. 2008 [cited 2023 Feb 26];31(8):1619–30. Available from: https://pubmed.ncbi.nlm.nih.gov/18971538/

38. John ASP, Kundu S, Pushpakumar S, Fordham M, Weber G, Mukhopadhyay M, et al. GYY4137, a Hydrogen Sulfide Donor Modulates miR194-Dependent Collagen Realignment in Diabetic Kidney. Scientific Reports 2017 7:1 [Internet]. 2017 Sep 7 [cited 2023 Feb 26];7(1):1–20. Available from: https://www.nature.com/articles/s41598-017-11256-3

39. Guan R, Wang J, Cai Z, Li Z, Wang L, Li Y, et al. Hydrogen sulfide attenuates cigarette smoke-induced airway remodeling by upregulating SIRT1 signaling pathway. Redox Biol. 2020 Jan 1;28:101356.

40. Chen Y, Yuan S, Cao Y, Kong G, Jiang F, Li Y, et al. Gasotransmitters: Potential Therapeutic Molecules of Fibrotic Diseases. Oxid Med Cell Longev. 2021;2021.

41. Sonke E, Verrydt M, Postenka CO, Pardhan S, Willie CJ, Mazzola CR, et al. Inhibition of endogenous hydrogen sulfide production in clear-cell renal cell carcinoma cell lines and xenografts restricts their growth, survival and angiogenic potential. Nitric Oxide [Internet]. 2015 Jun 26 [cited 2023 May 3];49:26–39. Available from: https://pubmed.ncbi.nlm.nih.gov/26068241/

42. Marutani E, Morita M, Hirai S, Kai S, Grange RMH, Miyazaki Y, et al. Sulfide catabolism ameliorates hypoxic brain injury. Nature Communications 2021 12:1 [Internet]. 2021 May 25 [cited 2023 May 3];12(1):1–19. Available from: https://www.nature.com/articles/s41467-021-23363-x

43. Huang Y, Wang G, Zhou Z, Tang Z, Zhang N, Zhu X, et al. Endogenous Hydrogen Sulfide Is an Important Factor in Maintaining Arterial Oxygen Saturation. Front Pharmacol. 2021 May 31;12:1366.

44. Forgan LG, McNeill BA, DeLeon E, Gao Y, Olson K. Effects of Hypoxia on Hydrogen Sulfide Production and Degradation Gene Expression Pathways. The FASEB Journal [Internet]. [cited 2023 May 3];31:700.9–700.9. Available from: https://onlinelibrary.wiley.com/doi/full/10.1096/fasebj.31.1_supplement.700.9

45. Uba T, Matsuo Y, Sumi C, Shoji T, Nishi K, Kusunoki M, et al. Polysulfide inhibits hypoxia-elicited hypoxia-inducible factor activation in a mitochondria-dependent manner. Mitochondrion. 2021 Jul 1;59:255–66.

46. Budde MW, Roth MB. Hydrogen sulfide increases hypoxia-inducible factor-1 activity independently of von Hippel-Lindau tumor suppressor-1 in C. elegans. Mol Biol Cell. 2010 Jan 1;21(1):212–7.

47. Wu B, Teng H, Zhang L, Li H, Li J, Wang L, et al. Interaction of Hydrogen Sulfide with Oxygen Sensing under Hypoxia. Oxid Med Cell Longev [Internet]. 2015 [cited 2023 May 3];2015. Available from: https://pubmed.ncbi.nlm.nih.gov/26078818/

48. Ellmers L, Templeton E, … APIJ of, 2020 undefined. Hydrogen sulfide treatment improves post-infarct remodeling and long-term cardiac function in CSE knockout and wild-type mice. mdpi.com [Internet]. [cited 2023 May 3]; Available from: https://www.mdpi.com/744314

49. Teuscher AC, Jongsma E, Davis MN, Statzer C, Gebauer JM, Naba A, et al. The in-silico characterization of the Caenorhabditis elegans matrisome and proposal of a novel collagen classification. Matrix Biol Plus. 2019 Feb;1:100001.

50. Choe KP, Strange K. Genome-wide RNAi screen and in vivo protein aggregation reporters identify degradation of damaged proteins as an essential hypertonic stress response. Am J Physiol Cell Physiol [Internet]. 2008 Dec [cited 2023 Feb 27];295(6):1488–98. Available from: www.ajpcell.org

51. Hine C, Harputlugil E, Zhang Y, Ruckenstuhl C, Lee BC, Brace L, et al. Endogenous hydrogen sulfide production is essential for dietary restriction benefits. Cell [Internet]. 2015 Jan 15 [cited 2023 Feb 27];160(1–2):132–44. Available from: https://pubmed.ncbi.nlm.nih.gov/25542313/

52. Qabazard B, Ahmed S, Li N, Arlt VM, Moore PK, Stürzenbaum SR. C. elegans aging is modulated by hydrogen sulfide and the sulfhydrylase/cysteine synthase cysl-2. PLoS One. 2013 Nov 8;8(11).

53. Qabazard B, Li L, Gruber J, Peh MT, Ng LF, Kumar SD, et al. Hydrogen sulfide is an endogenous regulator of aging in Caenorhabditis elegans. Antioxid Redox Signal [Internet]. 2014 Jun 1 [cited 2023 Feb 27];20(16):2621–30. Available from: https://pubmed.ncbi.nlm.nih.gov/24093496/

54. Ng LT, Ng LF, Tang RMY, Barardo D, Halliwell B, Moore PK, et al. Lifespan and healthspan benefits of exogenous H2S in C. elegans are independent from effects downstream of eat-2 mutation. npj Aging and Mechanisms of Disease 2020 6:1 [Internet]. 2020 Jun 10 [cited 2023 Feb 27];6(1):1–14. Available from: https://www.nature.com/articles/s41514-020-0044-8

55. Lohakul J, Jeayeng S, Chaiprasongsuk A, Torregrossa R, Wood ME, Saelim M, et al. Mitochondria-Targeted Hydrogen Sulfide Delivery Molecules Protect Against UVA-Induced Photoaging in Human Dermal Fibroblasts, and in Mouse Skin In Vivo. Antioxid Redox Signal [Internet]. 2022 Jun 1 [cited 2023 Apr 30];36(16–18):1268–88. Available from: https://pubmed.ncbi.nlm.nih.gov/34235951/

56. Rahimi M, Sohrabi S, Murphy CT. Novel elasticity measurements reveal C. elegans cuticle stiffens with age and in a long-lived mutant. Biophys J. 2022 Feb 15;121(4):515–24.

57. Pu YZ, Wan QL, Ding AJ, Luo HR, Wu GS. Quantitative proteomics analysis of Caenorhabditis elegans upon germ cell loss. J Proteomics. 2017 Mar 6;156:85–93.

58. Wei Y, Kenyon C. Roles for ROS and hydrogen sulfide in the longevity response to germline loss in Caenorhabditis elegans. Proc Natl Acad Sci U S A [Internet]. 2016 May 17 [cited 2023 Feb 27];113(20):E2832–41. Available from: https://www.pnas.org/doi/abs/10.1073/pnas.1524727113

59. Bolger AM, Lohse M, Usadel B. Trimmomatic: a flexible trimmer for Illumina sequence data. Bioinformatics. 2014 Aug 1;30(15):2114–20.

60. Dobin A, Davis CA, Schlesinger F, Drenkow J, Zaleski C, Jha S, et al. STAR: ultrafast universal RNA-seq aligner. Bioinformatics. 2013 Jan 1;29(1):15–21.

61. Anders S, Huber W. Differential expression analysis for sequence count data. Genome Biol. 2010 Oct 27;11(10):R106.

62. Love MI, Huber W, Anders S. Moderated estimation of fold change and dispersion for RNA-seq data with DESeq2. Genome Biol. 2014 Dec 5;15(12):550.

63. Ritchie ME, Phipson B, Wu D, Hu Y, Law CW, Shi W, et al. limma powers differential expression analyses for RNA-sequencing and microarray studies. Nucleic Acids Res. 2015 Apr 20;43(7):e47.

64. Adrian Alexa and Jörg Rahnenführer. topGO: Enrichment Analysis for Gene Ontology. R package version 2.46.0. 2021. topGO: Enrichment Analysis for Gene Ontology. R package version 2.46.0.

65. Alexa A, Rahnenführer J, Lengauer T. Improved scoring of functional groups from gene expression data by decorrelating GO graph structure. Bioinformatics. 2006 Jul 1;22(13):1600–7.

66. Raivo Kolde. https://CRAN.R-project.org/package=pheatmap. 2019. pheatmap: Pretty Heatmaps. R package version 1.0.12.

67. Kage-Nakadai E, Kobuna H, Kimura M, Gengyo-Ando K, Inoue T, Arai H, et al. Two very long chain fatty acid acyl-CoA synthetase genes, acs-20 and acs-22, have roles in the cuticle surface barrier in Caenorhabditis elegans. PLoS One. 2010 Jan 25;5(1):e8857.

68. Rieckher M, Tavernarakis N. Caenorhabditis elegans Microinjection. Bio Protoc. 2017;

69. Stroustrup N, Ulmschneider BE, Nash ZM, López-Moyado IF, Apfeld J, Fontana W. The Caenorhabditis elegans Lifespan Machine. Nat Methods. 2013 Jul 12;10(7):665–70.

