## Supplementary Information for "A VHL-1;HIF-1/SQRD1/COL-88 axis links extracellular matrix formation with longevity in *Caenorhabditis elegans*"

**Supporting Information**

**Supplementary Figure 1**

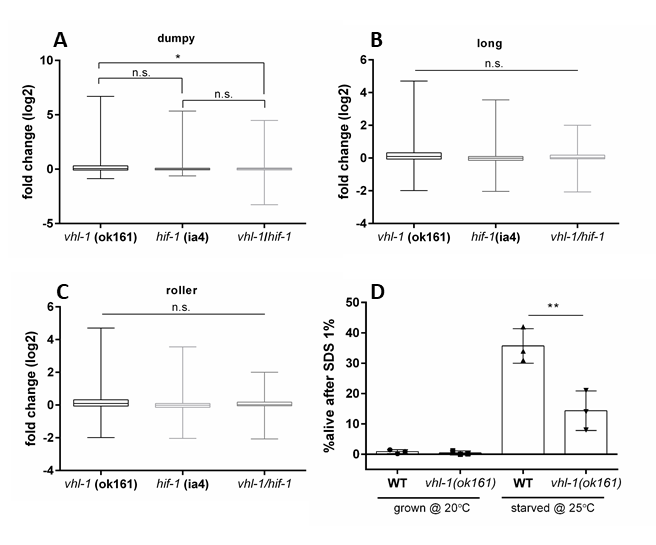

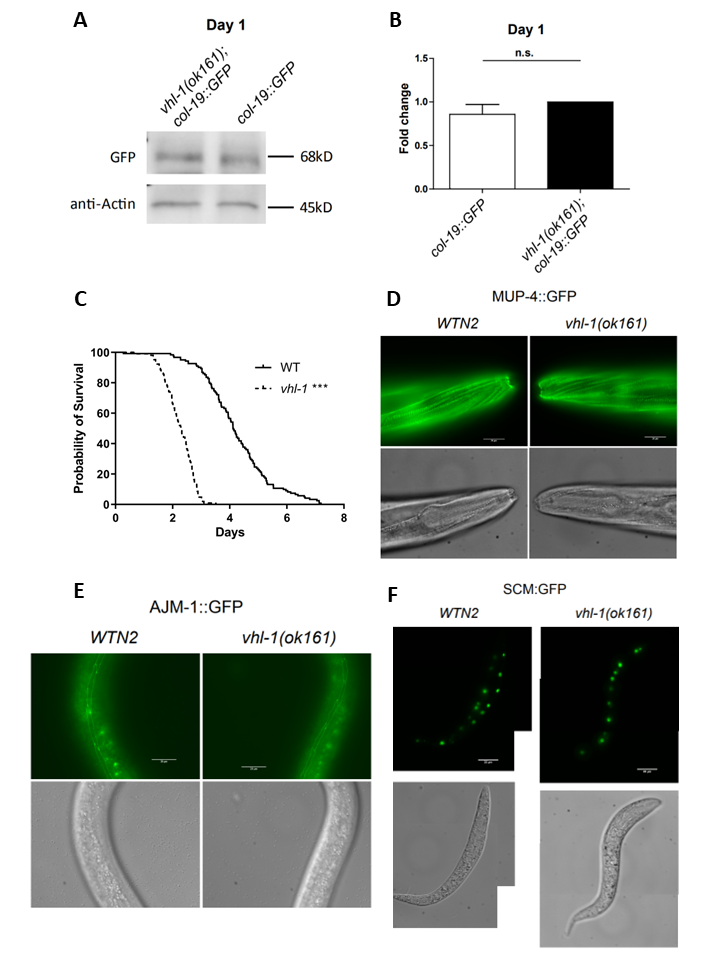
**Supplementary Figure 2.**

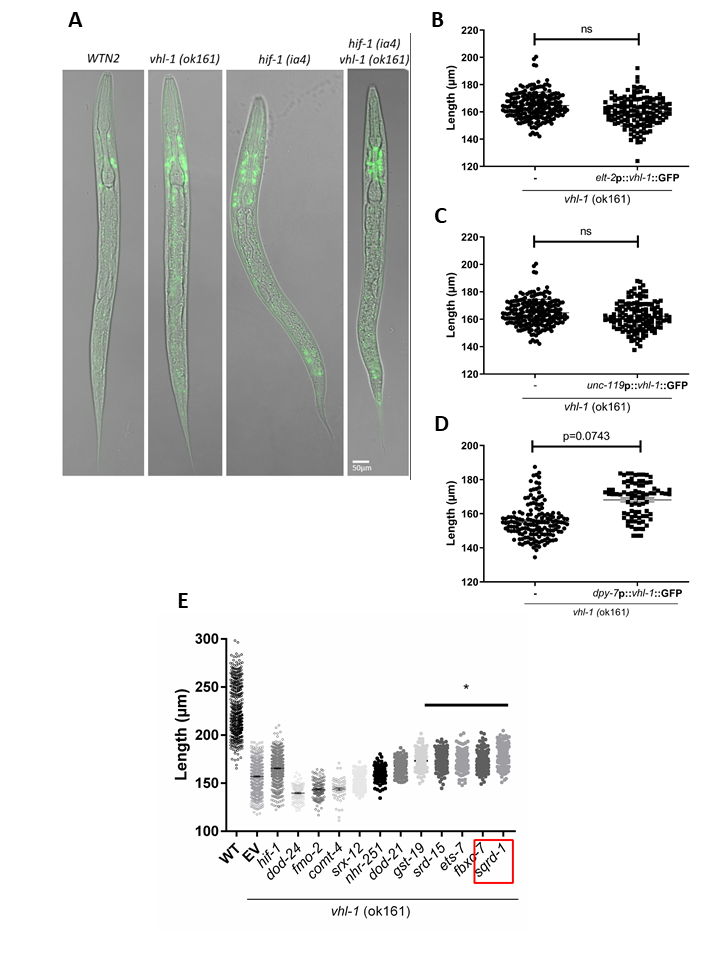
**Supplementary Figure 3.**

**Supplementary Figure 4.**

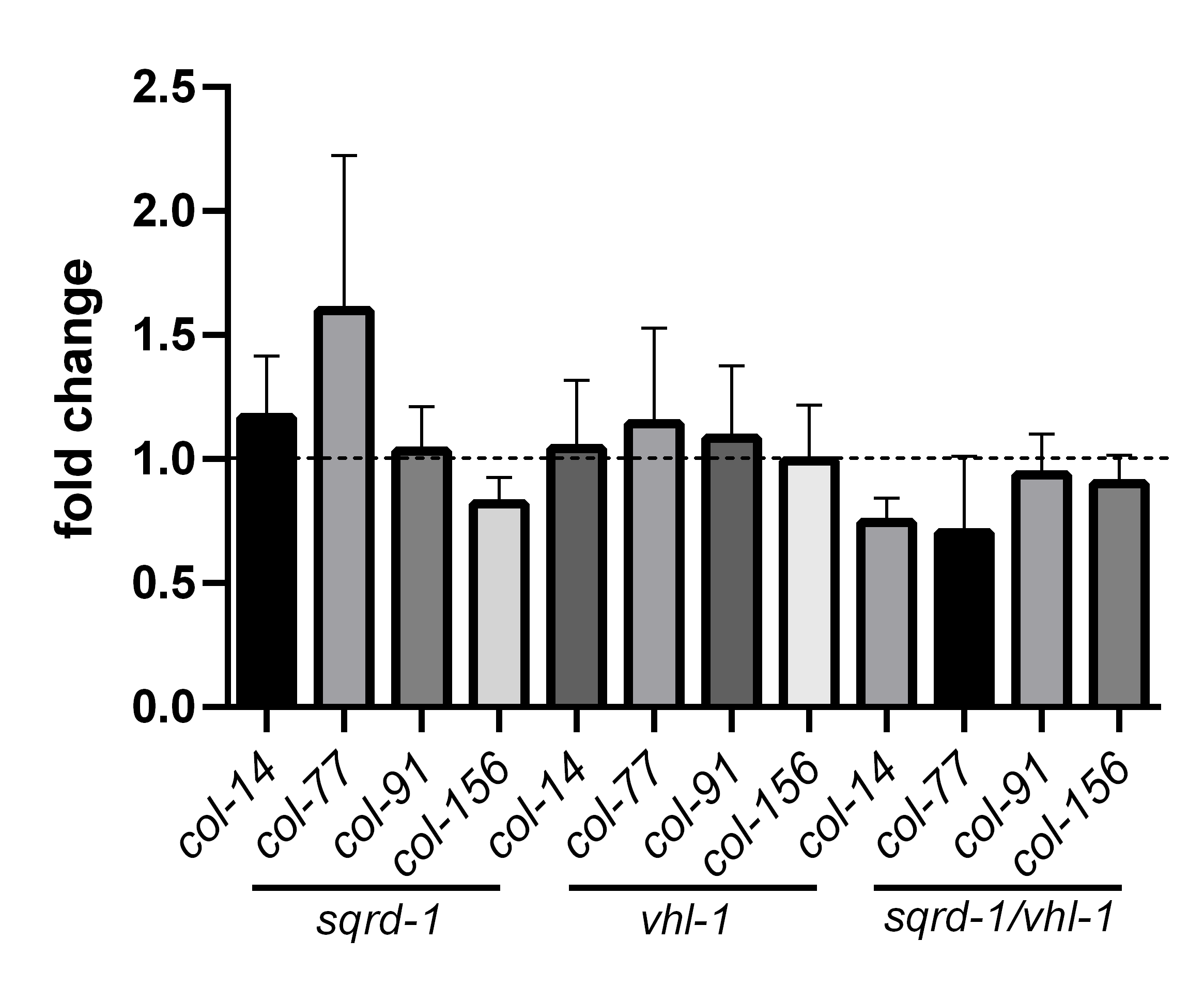

**Supplementary Table 1.**

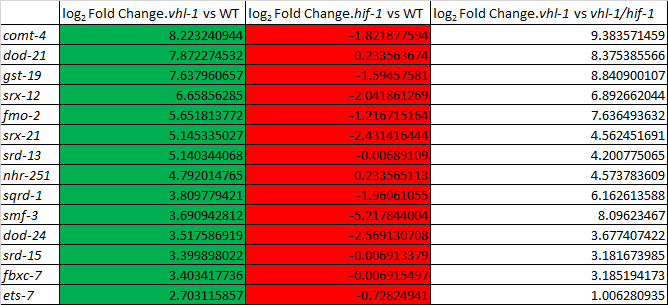

**Supplementary Table 2.**

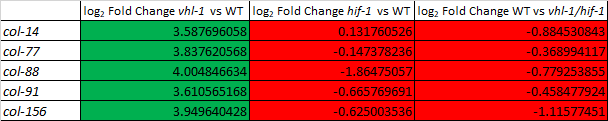

**Supplementary Table 3.**

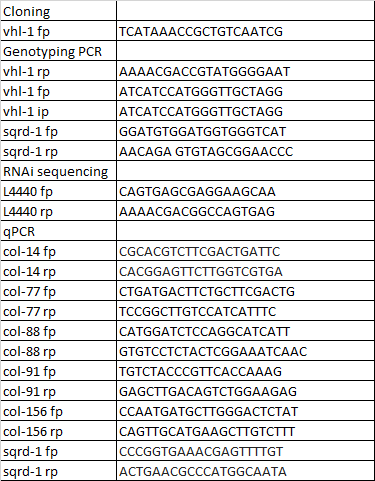
